# Spatiotemporal dynamics of nearshore fish communities in Casco Bay, Maine

**DOI:** 10.1101/2025.09.30.679511

**Authors:** Katie Lankowicz, Zach Whitener, Aaron Whitman, Samantha Bengs, Graham Sherwood, Courtney Swenson

## Abstract

Nearshore regions in the Gulf of Maine are highly variable habitats acted upon by processes operating on multiple spatial scales and levels of biological organization. As such, they facilitate the reproduction, growth, and migration of many fish species, and are particularly tied to the life histories of many forage fishes. Temperatures in the Gulf of Maine have rapidly increased in recent decades, which may be driving changes in nearshore ecosystems. Here, we use 11 years of summer beach seine survey data within Casco Bay, Maine, to illustrate temperature-related changes to community structure. Further, we use Atlantic herring (*Clupea harengus*) and Atlantic silversides (*Menidia menidia*) as focal species to describe the effects of temperature on individual growth rates and relative abundance. The progression of seasonal use patterns and relative abundance in the nearshore with seasonal warming is evident; species that have cooler preferred temperatures are caught less frequently in the nearshore regions in late summer, when temperatures are highest. Increased temperatures were associated with significantly higher silverside growth rates and community compositions dominated by silverside. Temperature alone did not explain interannual variation in herring growth rates or predict herring-dominated nearshore community composition, and there is evidence that density-dependence may be more important to herring population dynamics. Monitoring nearshore ecosystems could provide critical insight into the dynamics of species that use these areas to facilitate reproduction, growth, and migration, and could therefore be used to identify potential changes to Gulf of Maine community and trophic ecology.

## 1 Introduction

Nearshore regions of the ocean, via a unique combination of tidal mixing, freshwater input, shallow depths, complex habitats (e.g., rocky coastline or emergent macrophytes), and seasonal variation in temperatures, are important habitats for a wide array of aquatic species. The diverse mosaic of static (depth, substrate) and dynamic (tidal currents, wave action, temperature, salinity) environmental characteristics within these relatively small areas allows organisms to target one or more ecosystem services by selecting habitats at fine spatial and temporal scales. Nearshore regions are often used seasonally as spawning grounds (Fairchild et al., 2013) or nursery habitats (Stevenson et al., 2014) for juvenile life stages due to their high productivity and the protective effects of complex habitats and shallow and turbid waters against predation (Beck et al., 2001; Munsch et al., 2016). Nearshore regions also facilitate seasonal resource acquisition (Falke et al., 2024) and migratory pathways by both oceanodromous (Lazzari & Stevenson, 1992; Stevenson, 1989) and diadromous species (Saunders et al., 2006). The reliance of many commercially important species (e.g. herrings, groundfish, lobsters, bivalves) on nearshore regions for at least a portion of their life histories and the relative accessibility of these areas to human harvest conveys not only ecological importance, but also economic value.

Temperature is a critical factor for habitat selection, growth, and phenology for many aquatic species (Pörtner, 2002; Staudinger et al., 2019). Increases in ocean temperatures linked to climate change have been shown to affect habitat selection (Methratta & Link, 2007), distributions (Nye et al., 2009; Pershing et al., 2021), mortality and productivity (Le Bris et al., 2018; Pershing et al., 2015), and body sizes (Cheung et al., 2013; Sheridan & Bickford, 2011) of marine organisms. The response to increased temperatures can be species-specific and further affected by position along a latitudinal range, leading to varied impacts on species interactions and other aspects of community ecology. The North Atlantic Ocean, and the Gulf of Maine (hereafter, GoM) in particular, has experienced rapid warming in recent decades (Pershing et al., 2015, 2021). Basin-scale hydroclimatic changes have already been linked to changes in nearshore ecosystem use and population dynamics of iconic GoM species like American lobster and Atlantic cod (Le Bris et al., 2018; Pershing et al., 2015; Record et al., 2024). They may also be facilitating northward range expansions of typically mid-Atlantic species like black sea bass and blue crabs (Bell et al., 2015; Johnson, 2015; McBride et al., 2018). The GoM has relatively low biodiversity, which could limit the resilience of its ecosystems to biotic and abiotic changes (McMahan & Grabowski, 2019; Steneck & Wahle, 2013).

Regional warming is expected to continue at an above-average rate in the GoM as compared to other oceanic ecoregions (Saba et al., 2016). Temperature-linked disturbances to historic nearshore community structure and function may become more frequent, intense, and prolonged. Monitoring nearshore regions will be critical to predicting and understanding how increased temperatures may affect biological organization levels from individual species to the overall GoM ecosystem. In this study, we will relate changes in nearshore surface temperature to changes in the community structure, relative abundance, and summer growth rates of nearshore fishes commonly encountered in a beach seine survey, using Atlantic silverside (*Menidia menidia*; hereafter, silverside) and Atlantic herring (*Clupea harengus*; hereafter, herring) as focal species.

Atlantic silverside may be the only truly annual fish in the Northwest Atlantic, with little evidence of individuals reaching age 2 (Conover, 2024). Spawning occurs in shallow nearshore waters in the spring, and young-of-year migrate to deeper offshore waters in the winter to maintain a warmer thermal habitat. Its expected that less than 1% of the year class returns from this overwintering period on the inner continental shelf to recolonize shallow nearshore and estuarine habitats and spawn (Conover, 2024; Conover & Murawski, 1982). Silverside are distributed from southern Florida, USA to the northern Gulf of St. Lawrence, Canada, implying a diversity of genetic and phenotypic adaptations to a wide range of temperatures. Casco Bay, Maine, is in the northern half of this spatial range. The rapid generation time and ubiquity of silversides to much of the North American Atlantic coast have made them a useful tool to study the varied effects of temperature on sex determination, countergradient growth variation, spatial scale of local adaptation, and fisheries selectivity-induced evolution (see the works of David O. Conover cited within Conover (2024)). Silversides have demonstrated sensitivity to temperature; when reared in common garden median-temperature experiments, higher-latitude fishes have increased growth rates when compared to lower-latitude conspecifics (Conover & Present, 1990). This response is consistent with an expected temperature-mediated growth increase for a species that is not near its thermal maximum and indicates that increased temperatures in Casco Bay would result in higher growth rates for local silversides.

Atlantic herring are distributed on both sides of the Atlantic, with the Gulf of Maine-Georges Bank complex representing the southernmost spawning population in the western Atlantic (Stevenson & Scott, 2005). Spawning occurs in the late summer to fall, with peak spawning occurring in October (Graham, Davis, et al., 1972). Larvae are dispersed into coastal zones where they overwinter (Graham, Chenoweth, et al., 1972; Graham & Townsend, 1985) and grow rapidly in the following spring and summer (Anthony, 1972; Lough et al., 1982). Until they reach sexual maturity around ages 3-4 (Chamberland et al., 2022; OBrien et al., 1993), herring will annually move offshore or to deeper bays to overwinter in more suitable thermal habitats (Boyar, 1968) and generally remain close to these overwintering grounds (Creaser & Libby, 1986, 1988). Adults are oceanodromous and will make seasonal migrations from more northerly spawning grounds to more southerly overwintering grounds. In the Northwest Atlantic, this overall distribution ranges from South Carolina, USA (southern extent, winter distribution) to Labrador, Canada (northern extent, summer distribution). Increased temperatures have been found to be associated with decreased growth rate (Sswat et al., 2018). A recent synoptic study of all Northwest Atlantic herring populations indicates that there is a consistent cross-population response of smaller body size with increased sea surface temperatures, and herring that spawn and overwinter in the warmer (southern) edge of the population range have the strongest response (BeaudrySylvestre et al., 2024). It has been difficult to definitively link warming ocean temperatures to recent declines in spawning stock biomass of US herring stocks, likely due to confounding effects of historical exploitation (Pershing et al., 2021) and the effects of density-dependence (Becker et al., 2020). Despite this, temperature has been shown to affect recruitment and phenology in Canadian stocks in the Bay of Fundy and Scotian Shelf (Boyce et al., 2021). Additionally, several studies predict a decrease in thermally suitable habitat and an associated decrease in abundance for herring in the GoM by 2050 (Allyn et al., 2020; Kleisner et al., 2017).

Both silverside and herring are highly abundant in the nearshore regions of the GoM in the summer. However, their differing life history strategies and the relative position of the GoM to their overall distributions (northern half of range for silverside, southern half of range for herring), could lead to opposing responses to increasing temperatures. Increases in nearshore GoM surface temperatures may be advantageous to silverside growth and reproduction, while conversely altering patterns of nearshore habitat use and reducing the growth of herring. In this study, we use 11 years of summer beach seine data collected in Casco Bay, Maine to explore the effect of nearshore temperature on the summer growth rates and relative abundances of herring and silverside. We also use these data to examine changes in spatiotemporal community structure. Seasonal warming and the linked phenology of nearshore species should result in juvenile stages of spring-spawning oceanodromous and diadromous fishes dominating species assemblages in the early summer. Alternatively, summer-spawning oceanodromous fishes and nearshore residents should dominate in the late summer. Average annual community composition across all of Casco Bay may shift from herring-dominated in cooler years to silverside-dominated in recent, warmer years.

## 2 Methods

### 2.1 Field data collection

Since 2014, the Gulf of Maine Research Institute has conducted summer beach seines at 12 sites within the western half of Casco Bay as part of the Casco Bay Aquatic Systems Survey (CBASS) program. Sites range from just north of the Presumpscot River mouth to just north of Trundy Point, Cape Elizabeth (Fig. 1), resulting in samples arranged along a gradient of salinity. Bottom substrate type varies across sites from fine-grained mud at the sites within the Presumpscot River to coarse-grained gravel at sites along the Cape Elizabeth coast. Sites were sampled at approximately 2-week intervals from early summer through early fall, with scattered omissions or delays due to weather or mechanical issues.

**Figure 1:**
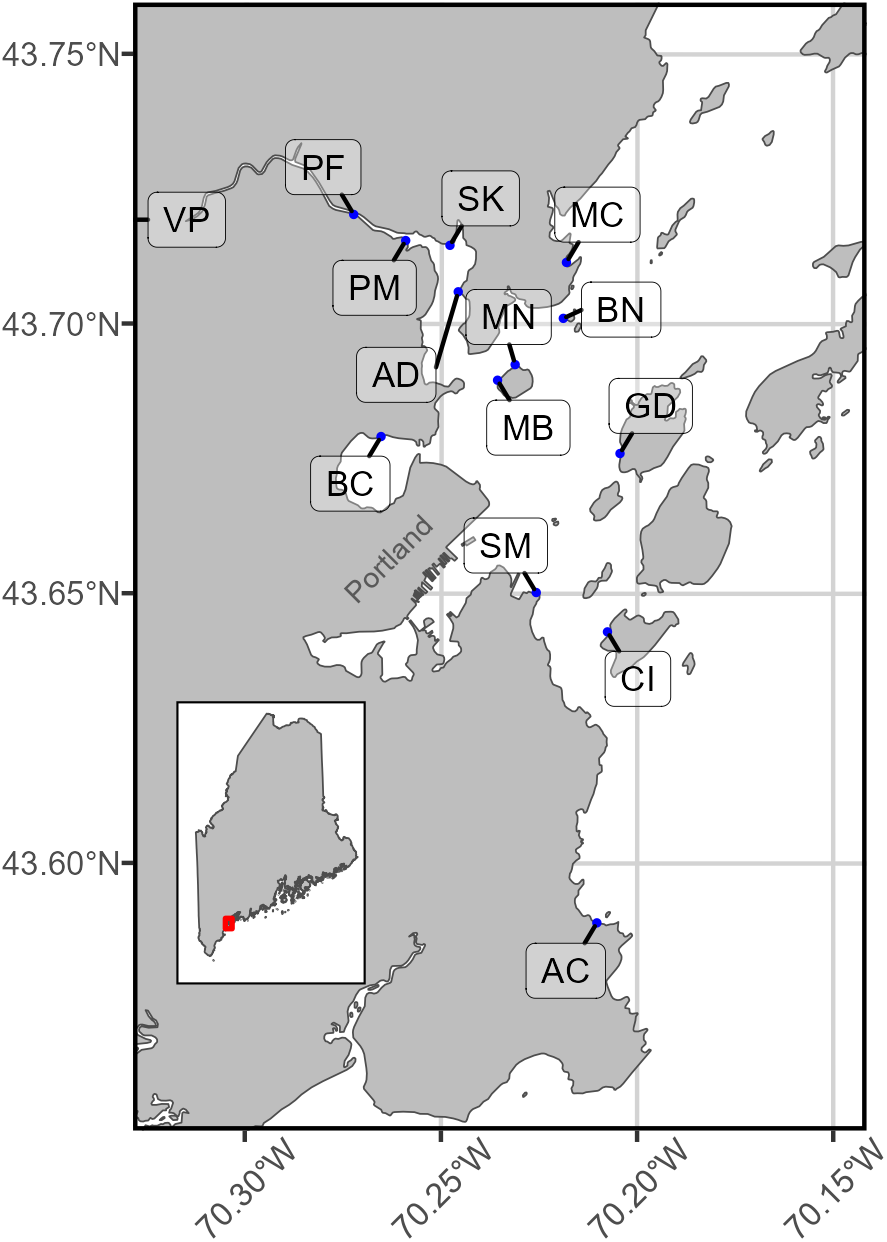
Location of sampling sites along the western coast of Casco Bay. Inset shows the state of Maine and relative position (red border) of the sampling area. Sites are, from north to south, Presumpscot Moorings (PM), Skitterygusset (SK), Mussel Cove (MC), Audubon (AD), Brothers Island North (BN), Mackworth Island North (MN), Mackworth Island Beach (MB), Back Cove (BC), Great Diamond Island (GD), Southern Maine Community College (SM), Cushing Island (CI), and Alewife Cove (AC). The city of Portland is also labeled for context.

All samples were collected using a 45.7 m long, 2.4 m tall seine built by Memphis Net and Twine of 4.8 mm knotless delta-style nylon mesh with a central 2.4m by 2.4m by 2.4m bag. The bottom line was weighted with lead sinkers and the top line was fitted with buoyant Spongex floats. The seine was connected to a bridle and 2m of spare line on both ends. Seines were deployed with the help of a skiff. For each sampling event, one end of the seine was held on the beach while the net was flaked out from the bow of the skiff as it backed off the beach. The skiff was moved to extend the seine to the near edge of the bag, then pivoted to return to shore. The remaining length of the seine was flaked out until the skiff landed back on the beach. This resulted in a generally U-shaped deployment of the seine with approximately 20 m between the ends on the beach. Tidal stage was not a criterion for sampling, but tidal information was recorded and seines were set so that the deepest point sampled was between 0.5 and 2.2 m. Once both ends of the seine were on the beach, one person on each end hand hauled the net in. The bag was left partially in the water to limit specimen mortality. The contents of the bag were sorted by species and enumerated. The first 25 individuals of each species were measured to the nearest mm and species with easily identifiable sexually dimorphic features (i.e., crabs) were sexed. All organisms were released immediately after they were counted and/or measured. After completion of seine operations, surface temperature and salinity were measured using a YSI multiparameter sonde, and general weather conditions were recorded.

Data cleaning and processing were accomplished before beginning any statistical analyses. Minimal sampling occurred in 2019 and therefore 2019 data are excluded from all analyses. The most consistently sampled period each year was between weeks 24 and 39 (early June through mid-September). Sporadic data outside this period were removed. Data from observations that noted an issue with the setting of the seine (water depth below the minimum standard, net coming in tangled, etc.) were also excluded. Of 707 initial seine sampling records, 659 remained after data cleaning.

### 2.2 Nearshore surface temperature anomaly

Temperature data from each sampling event characterizes instantaneous local conditions but cannot characterize long-term changes in nearshore surface temperature. To create a more synoptic index of nearshore surface temperature, we used data from the NOAA Portland Harbor tide gauge (Station ID: 8418150). Although this station records the temperature of only one point within Casco Bay, it is the most complete dataset available for historical nearshore water temperature information. The station has reliably recorded surface water temperature at 6-minute intervals since August 2002 with minimal disruption. Wider spatial scale data sources, such as NOAAs Optimum Interpolated Sea Surface Temperature (OISST) product, do not predict the temperature of the extreme nearshore shallows sampled in this study, and thus may not create a precise or accurate index of temperatures in these areas.

We extracted temperature data from January 2003 through December 2024, cleaned to remove ecologically unlikely values (sudden spikes indicative of the instrument being removed from water, or temperatures above 30°C), and calculated the mean daily temperature. The climatological reference period (CRP) was defined as January 2003 through December 2020. Standard practice, as described by the NOAA National Centers for Environmental Information, is to use a defined 30-year period to calculate climatological norms. The current United States national CRP is 1991-2020. The Portland Harbor tide gauge does not have an instrument record extending back this far, so we chose to use the longest uniform period in the instrument record as our CRP. A Generalized Additive Model (GAM) was used to estimate daily mean temperature over the CRP. Day of year and year were used as the explanatory variables. The mgcv package (S. N. Wood, 2004) was used to fit the model. Predictions of mean temperature per day of year, excluding the effects of individual years, were extracted.

Daily temperature anomalies within the sampling period (2014-2024) were calculated as the difference between the mean daily temperature at the Portland Harbor tide gauge and the predicted daily temperature of the CRP. The temporal frame of reference was shifted so that day 1 of each modeled year was December 1st. This was done to best align the categorization of temperature anomalies with seasonal temperature changes that would impact summertime ecosystem dynamics; winter (December-February) and spring (March-May) temperatures are critical for the timing and success of reproduction and growth for many GoM species. Mean annual temperature anomalies were calculated according to this frame of reference (December 1st through the following November 30th). Years with mean temperature anomalies below the CRP average were categorized as cooler, and years with mean temperature anomalies above the CRP average were categorized as warmer.

### 2.3 Statistical analyses

#### 2.3.1 Weekly growth rates

Growth rate fluctuates with age, so it is necessary to separate fish captured in summer beach seine operations into discrete year classes. When direct measurements of age are not available, statistical length-frequency models can be used to estimate age. This process is effective for temperate fishes with short, distinct spawning seasons (Macdonald & Pitcher, 1979). The relationship of length to age becomes more difficult to detect as fish age and growth slows or ceases (Maunder et al., 2018). However, both focal species (herring and silverside) can likely be aged using this approach. Both species have seasonal growth patterns. The short lifespan and simple age structure of silverside should facilitate the detection of distinct ages by size. Though older and larger herring may no longer have distinct distributions of size at age, all herring individuals caught in our seine sampling efforts were 14-139 mm long. When comparing to the expected size and age of herring at maturation (275 mm at ages 3-4; (Boyar, 1968)) it is clear that these are all younger individuals which should have detectable relationships of length to age.

Next, we identified and selected probable age-0 individuals of both species for the calculation of weekly growth rates. This begins with an assumption that the lengths of sampled conspecifics in the same year class at the same time come from an identifiable distribution. Commonly, a normal distribution is assumed (Macdonald & Pitcher, 1979; Zhou et al., 2022). It is therefore possible to identify age groups by their unique modal lengths along a cumulative length distribution (Macdonald & Pitcher, 1979). To confirm the presence of age-0 fish, the weekly length distributions of silversides and herring were visualized (Fig. 2) and compared to published values of length at age. Juvenile herring grow to lengths of 90-125 mm by the end of their first year (Anthony, 1972). The size range of herring caught in our beach seines indicates they were mostly age-0 fish spawned the previous fall (mean 64 mm, SD 13 mm). Silverside spawn in the spring, with juveniles reaching 90-100 mm before growth ceases in the late fall (Conover & Ross, 1982). Silverside weekly average lengths were consistently around 100 mm in the first third of the sampling season but rapidly decreased to a low of 67 mm in week 31. This indicates that mostly age-1+ silversides were caught early in the season, but recently-spawned silversides recruit to the seine and dominate catch by late July. This is consistent with phenological patterns and growth rates reported in the literature (Conover & Present, 1990; Conover & Ross, 1982; Gao & Munch, 2013).

**Figure 2:**
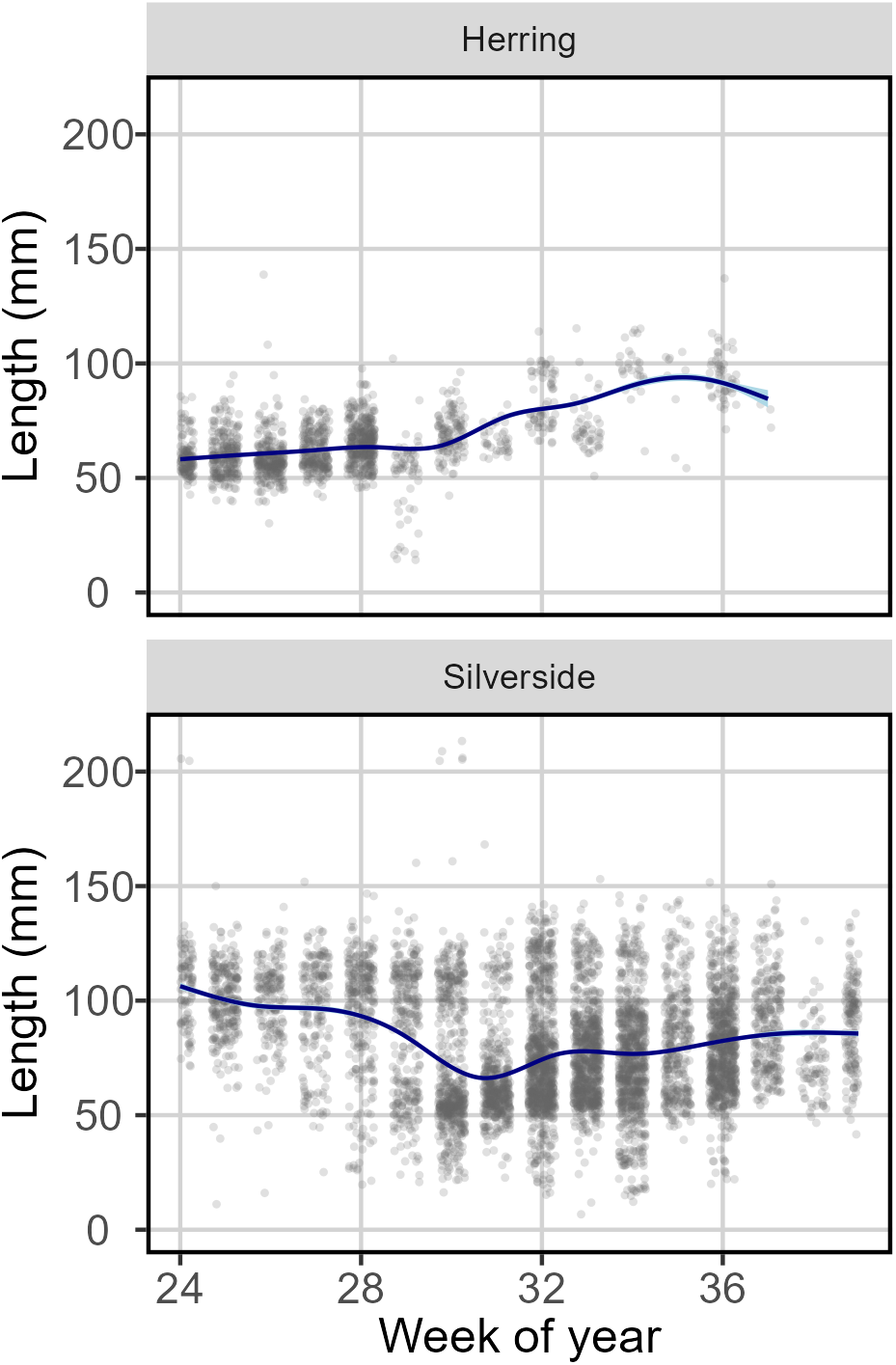
Weekly length distributions (in mm) for herring (top panel) and silverside (bottom panel) aggregated across all sampled years. Blue lines indicate locally estimated scatterplot smoothing (LOESS) regressions to illustrate changes average weekly lengths.

The following analyses were carried out separately for each year of data collection, as we expected interannual variation in spawning timing and growth. Bayesian inference through the R package LaplacesDemon (Statisticat LLC, 2021) was used to estimate the number of length distribution modes within each week. This was accomplished through the Modes function, which is a deterministic function that differences the kernel density of a continuous variable and reports n modes equal to half the number of changes in direction. This allows for the statistical determination of the number of age groups, as opposed to traditional length frequency analysis methods that require either a priori knowledge or graphical analysis to identify age structure (Fournier et al., 1990; Petersen, 1891; Zhou et al., 2022). If the assumption that the lengths of each age group come from a normal distribution holds, the location of each mode would represent the mean length of fish belonging to that group. The location of modes was estimated using a mixed modeling approach via the mixtools R package (Benaglia et al., 2009). An expectation-maximization (EM) algorithm for mixtures of univariate normals applied to the number of modes identified in the previous step was applied. A 95% confidence interval was estimated as 2 standard deviations around each identified mode and stood in for the estimated size range of each age group.

This approach to length-frequency analysis is similar to established software and methods (Gayanilo et al., 1996; Mildenberger et al., 2017; Zhou et al., 2022). Like these other approaches, an expert is required to apply informative priors and make decisions about the validity of the identified age groups. We developed a set of rules to exclude biologically invalid or improbable estimates of age-specific size range. For the former, this included length estimates with minimum lengths below 0mm or maximum lengths above the literature-derived maximum asymptotic body length. For the latter, this included length estimates where the range of possible lengths exceeded 40% of the value of the maximum asymptotic body length. It is unlikely that fish of the same age and captured in the same location would have such extreme variation in achieved lengths. We also merged groups with more than 50% overlap in estimated size confidence intervals, as this is not enough separation to identify distinct age groups. From the remaining data, we identified the first week each summer in which probable age-0 fish were identified, then tracked the growth of that cohort through the following weeks.

Next, the lower and upper limits of the length range of the age-0 cohort for every week in the sampling period were estimated. This was necessary to create a continuous time series of estimated lengths; data volume limitations prevented the Bayesian and mixed modeling approach from identifying an age-0 cohort in some weeks. Lifetime fish growth is usually modeled using the von Bertalanffy equation, which is a specialized case of logistic growth. Beach seine operations took place over 15 weeks each summer, with age-0 cohorts for both focal species persisting in the area for only a few weeks of that time. Rapid growth over this relatively short period supports estimating the observed growth rate as a linear relationship. Therefore, linear models were fit to the weekly upper and lower limits of length ranges as estimated in the previous step. Fish with lengths within the modeled limits of each week were assigned to the age-0 cohort. Fish not assigned to the age-0 cohort were removed from further growth analyses.

The weekly length distributions of both age-0 silversides and age-0 herring in each year often exhibited higher variance in later weeks as compared to earlier weeks. Weighted least squares regressions were used to control for this heteroskedasticity. In these models, the lengths of the selected fishes were the response variable and the numeric week of the year was the explanatory variable. The weekly growth rate of the age-0 cohort of each species in each year was therefore defined as the slope of the regression fit to the weekly age-0 cohort length frequency distribution data. Two-sided Welchs t-tests were used to compare weekly growth rates within each species between years with an annual temperature above the CRP average (warmer years) and those with an annual average temperature below the CRP average (cooler years).

#### 2.3.2 Effects on catch

Generalized additive models were used to model changes in herring and silverside catch along environmental gradients and through time. Catch data for herring and silverside was filtered to only include observations where the time of the observation and the surface temperature were recorded. It is important to note that, unlike the growth analysis, these GAMs use all catch data and do not focus on only age-0 fishes. GAMs were fit with a negative binomial error distribution family and a log-link function due to the zero-inflated and overdispersed nature of the catch data. Thin-plate regression splines were fit to all continuous numeric variables and interaction terms.

Environmental variables tested as explanatory variables included surface temperature (°C), substrate type, weather conditions, tidal state (rising or falling), and water level (m) as compared to the Mean Lower Low Water (MLLW) datum at the time and location of each seine haul. Surface temperature data were collected immediately after each seine. Though salinity data were also collected, there were several long gaps in data availability from sensor failures. Thus, salinity could not be included as an environmental variable. The substrate at each sampling site was categorized as mud, sand, or sand/gravel mixture according to an estimate of average grain size within the sampled area. Weather conditions were categorized into sunny, partly cloudy, overcast, or rainy. Both substrate type and weather conditions were included as fixed-effect factor variables. Tidal data were extracted from the nearby Portland Harbor tide gauge. Water level data were extracted at the native 6-minute frequency, then interpolated and smoothed to a 1-minute frequency for every summer when sampling occurred. The timestamp of each seine set was then used to determine the tidal state (rising or falling; fixed-effect factor variable) and approximate water level as compared to the MLLW datum. Though the gauge measurements are specific to Portland Harbor, the time offsets for high and low tides at subordinate NOAA tidal prediction stations near the edges of Casco Bay are between 1 and 2 minutes and tidal height offsets are between 0 and 0.3 m. All our sampled sites are relatively close to Portland Harbor and would likely have temporal and tidal height offsets within this range, which is small enough that we would not likely be able to detect any ecological effects.

Temporal variables included year and numeric week of year. Year was treated as a fixed-effect factor variable so that the effect of each year on catch could be estimated independently, as is most appropriate for generating indices of relative abundance. Week was incorporated as an interaction term with surface temperature. This was done to account for the effect of seasonal warming on the phenology of the focal species, particularly herring. If warming signals and temperature maxima are earlier each year, herring use of the nearshore area may end earlier, and herring catch may decrease. The interaction between week of year and surface temperature could also highlight age-specific temperature preferences, particularly for silversides.

A random effect for each site was also included as a model term. Though we were not interested in characterizing the difference in catch between sites, including this term was necessary to account for the temporal autocorrelation introduced by the repeated measures sampling design. Site was included as a random rather than a fixed effect because it is likely that the catch of fish at any location is a function of the interaction of multiple local conditions, including those beyond what was explicitly included in the model (e.g., bathymetric features or shoreline shapes that influence currents, concentration of planktonic prey items, salinity). The sites in our study design can be thought of as random samples along the gradients of those unmeasured environmental conditions. Term selection was conducted according to the double penalty, or null space penalization, approach (Marra & Wood, 2011). This enables shrinkage for all smoothed terms and essentially penalizes insignificant terms to 0 through shrinking function components both in the range space and in the null space.

#### 2.3.3 Spatiotemporal community structure

The relative proportion of observed species is variable across the different sampling sites within Casco Bay, as each site has a unique set of static and dynamic environmental characteristics that can influence species-specific habitat selection patterns. Seasonal warming can trigger shifts in habitat use patterns, especially for migratory fishes seeking a preferred thermal range. To get a fine-scale assessment of nearshore community structure, the catch per unit effort (CPUE) of every encountered species was calculated at each site within three seasonal periods. Interannual variability was not considered at this stage, so results were pooled across all years. The seasonal periods were defined as early summer (weeks 24-28 of each year), mid summer (weeks 29-33), and late summer (weeks 34-39). CPUE per seasonal period for each species was calculated as the number of individuals caught divided by the number of seine hauls. The data were filtered so that only species that were caught in more than 1% of all seine hauls and across 8 of 10 sampling years remained. This approach was used to limit the influence of uncommon and possibly misidentified species. Encounter rates were used as thresholds rather than total abundance to avoid bias towards schooling or shoaling species. After this process, 10 species remained of 42 total species encountered.

Community composition analysis was conducted using nonmetric multidimensional scaling (NMDS), which ordinates data without assuming linear relationships or distance metric properties (Clarke, 1993). This approach was applied to Bray-Curtis similarity matrices. Because catch was standardized by effort in the previous step, no transformation was applied. The goodness of fit of the data across the ordination axes was indexed by the stress coefficient, which represents distortion from reducing multidimensional data to fewer dimensions. If the calculated stress coefficient was less than 0.2, the NMDS plot was deemed an acceptable representation of the data (Clarke, 1993; Field et al., 1982).

After NMDS, k-means clustering was used to identify groups with similar community structures. This is a non-hierarchical method used to minimize the distance between data points and the center of an assigned cluster. The silhouette and total within sum of squares (WSS) methods were used to identify the most appropriate number of clusters. We did not assign data points a priori to clusters by seasonal period or location within the study area, as we expected both spatial location and seasonal warming to affect community composition and did not want to assume the nature of that interaction. A permutation-based, one-way Analysis of Similarities (ANOSIM) test was used to identify whether community composition was statistically significantly different between clusters (Clarke, 1993). This was followed by a Similarity Percentages (SIMPER) test, which was used to identify the species that contributed the most to the dissimilarity between identified clusters community composition (Clarke, 1993). All statistical tests except k-means clustering were performed using the vegan R package, version 2.6-8 (Oksanen et al., 2024).

Interannual differences in community structure, driven by larger-scale and longer-term processes than those encompassed by the previous analysis, were also of interest. We applied the same sequence of testsNMDS, k-means clustering, ANOSIM, and SIMPERon species CPUE when pooled across all sampling sites and seasonal periods per year.

## 3 Results

### 3.1 Species caught

In total, 659 seine hauls across 132 unique sampling days met data collection standards and were included in our analyses (Table 1). The total catch was 159,590 individuals across 42 species. The 10 species and/or genera that met encounter thresholds to be included in community composition analyses were, in order of increasing encounter percentage, bluefish (*Pomatomus saltatrix*), northern pipefish (*Syngnathus fuscus*), American sand lance (*Ammodytes americanus*), Atlantic tomcod (*Microgadus tomcod*), Atlantic herring (*Clupea harengus*), river herring (*Alosa spp*.), mummichog (*Fundulus heteroclitus*), winter flounder (*Pseudopleuronectes americanus*), Atlantic silverside (*Menidia menidia*), and green crab (*Carsinus maenas*). Note that river herring is an umbrella term for both alewife (*Alosa pseudoharengus*) and blueback herring (*Alosa aestivalis*). We did not differentiate between the two, as they are difficult to identify without lethal sampling. Combined, these 10 commonly-encountered species comprised over 99% of all individuals caught. Atlantic silverside was the most abundant species at nearly 42% of total catch and was encountered in 56% of all seine hauls. Atlantic herring were also highly abundant (nearly 35% of total catch), but were encountered in only 15% of all seine hauls. Both species were encountered in every year of the sampling period. Green crabs had the widest range of spatiotemporal distribution and were encountered in 68% of all seine hauls. With few exceptions, they were detected at least once at every site in every year. Green crabs are the only non-teleost included in the analyses; other benthic invertebrates like horseshoe crabs (Limulus polyphemus) and various species of shrimp and hermit crabs were occasionally encountered but not enumerated.

**Table 1:**
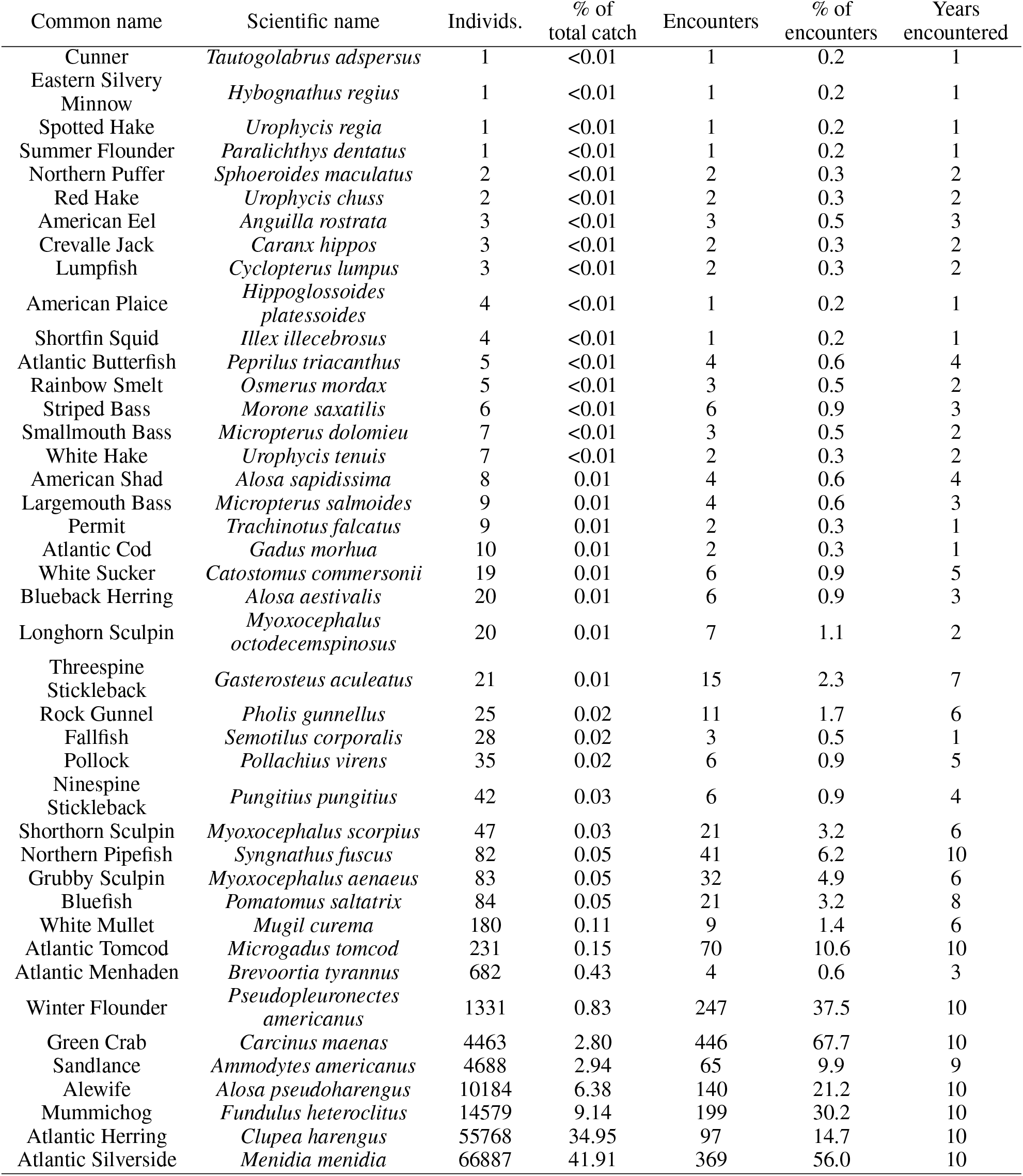
Summary of seine catch data of 11 year time series. Species are arranged by number of individuals caught, number of seine hauls where at least one individual of the species was encountered (Encounters), and number of years where at least one individual of the species was encountered (Years encountered). The percentage of total catch (number caught divided by 159,590 total organisms enumerated) and percentage of encounters (number of seine hauls seen divided by 659 total seine hauls) are also calculated.

| Common name | Scientific name | Individs. | % of total catch | Encounters | % of encounters | Years encountered |
| --- | --- | --- | --- | --- | --- | --- |
| Cunner | *Tautogolabrus adspersus* | 1 | <0.01 | 1 | 0.2 | 1 |
| Eastern Silvery Minnow | *Hybognathus regius* | 1 | <0.01 | 1 | 0.2 | 1 |
| Spotted Hake | *Urophycis regia* | 1 | <0.01 | 1 | 0.2 | 1 |
| Summer Flounder | *Paralichthys dentatus* | 1 | <0.01 | 1 | 0.2 | 1 |
| Northern Puffer | *Sphoeroides maculatus* | 2 | <0.01 | 2 | 0.3 | 2 |
| Red Hake | *Urophycis chuss* | 2 | <0.01 | 2 | 0.3 | 2 |
| American Eel | *Anguilla rostrata* | 3 | <0.01 | 3 | 0.5 | 3 |
| Crevalle Jack | *Caranx hippos* | 3 | <0.01 | 2 | 0.3 | 2 |
| Lumpfish | *Cyclopterus lumpus* | 3 | <0.01 | 2 | 0.3 | 2 |
| American Plaice | *Hippoglossoides platessoides* | 4 | <0.01 | 1 | 0.2 | 1 |
| Shortfin Squid | *Illex illecebrosus* | 4 | <0.01 | 1 | 0.2 | 1 |
| Atlantic Butterfish | *Peprilus triacanthus* | 5 | <0.01 | 4 | 0.6 | 4 |
| Rainbow Smelt | *Osmerus mordax* | 5 | <0.01 | 3 | 0.5 | 2 |
| Striped Bass | *Morone saxatilis* | 6 | <0.01 | 6 | 0.9 | 3 |
| Smallmouth Bass | *Micropterus dolomieu* | 7 | <0.01 | 3 | 0.5 | 2 |
| White Hake | *Urophycis tenuis* | 7 | <0.01 | 2 | 0.3 | 2 |
| American Shad | *Alosa sapidissima* | 8 | 0.01 | 4 | 0.6 | 4 |
| Largemouth Bass | *Micropterus salmoides* | 9 | 0.01 | 4 | 0.6 | 3 |
| Permit | *Trachinotus falcatus* | 9 | 0.01 | 2 | 0.3 | 1 |
| Atlantic Cod | *Gadus morhua* | 10 | 0.01 | 2 | 0.3 | 1 |
| White Sucker | *Catostomus commersonii* | 19 | 0.01 | 6 | 0.9 | 5 |
| Blueback Herring | *Alosa aestivalis* | 20 | 0.01 | 6 | 0.9 | 3 |
| Longhorn Sculpin | *Myoxocephalus octodecemspinosus* | 20 | 0.01 | 7 | 1.1 | 2 |
| Threespine Stickleback | *Gasterosteus aculeatus* | 21 | 0.01 | 15 | 2.3 | 7 |
| Rock Gunnel | *Pholis gunnellus* | 25 | 0.02 | 11 | 1.7 | 6 |
| Fallfish | *Semotilus corporalis* | 28 | 0.02 | 3 | 0.5 | 1 |
| Pollock | *Pollachius virens* | 35 | 0.02 | 6 | 0.9 | 5 |
| Ninespine Stickleback | *Pungitius pungitius* | 42 | 0.03 | 6 | 0.9 | 4 |
| Shorthorn Sculpin | *Myoxocephalus scorpius* | 47 | 0.03 | 21 | 3.2 | 6 |
| Northern Pipefish | *Syngnathus fuscus* | 82 | 0.05 | 41 | 6.2 | 10 |
| Grubby Sculpin | *Myoxocephalus aeneus* | 83 | 0.05 | 32 | 4.9 | 6 |
| Bluefish | *Pomatomus saltatrix* | 84 | 0.05 | 21 | 3.2 | 8 |
| White Mullet | *Mugil curema* | 180 | 0.11 | 9 | 1.4 | 6 |
| Atlantic Tomcod | *Microgadus tomcod* | 231 | 0.15 | 70 | 10.6 | 10 |
| Atlantic Menhaden | *Brevoortia tyrannus* | 682 | 0.43 | 4 | 0.6 | 3 |
| Winter Flounder | *Pseudopleuronectes americanus* | 1331 | 0.83 | 247 | 37.5 | 10 |
| Green Crab | *Carcinus maenas* | 4463 | 2.80 | 446 | 67.7 | 10 |
| Sandlance | *Ammodytes americanus* | 4688 | 2.94 | 65 | 9.9 | 9 |
| Alewife | *Alosa pseudoharengus* | 10184 | 6.38 | 140 | 21.2 | 10 |
| Mummichog | *Fundulus heteroclitus* | 14579 | 9.14 | 199 | 30.2 | 10 |
| Atlantic Herring | *Clupea harengus* | 55768 | 34.95 | 97 | 14.7 | 10 |
| Atlantic Silverside | *Menidia menidia* | 66887 | 41.91 | 369 | 56.0 | 10 |

The remaining 32 species, which combined made up less than 1% of total individuals caught, were mainly juvenile fishes native to the Gulf of Maine or connecting freshwater systems but also included permit (*Trachinotus falcatus*), crevalle jack (*Caranx hippos*), white mullet (*Mugil curema*), and summer flounder (*Paralychthis dentatus*). Permit and crevalle jack are among an assemblage of tropical or subtropical species that typically reside south of Cape Cod, but may be advected northward into the Gulf of Maine by eddies or warm core rings of the Gulf Stream Current (Hare et al., 2002; ONeill et al., 2025; A. J. M. Wood et al., 2009). Species within this group are colloquially known as Gulf Stream Orphans (gsoproject.org, see (ONeill et al., 2025)). Though not officially recognized as part of this group, white mullet and other Mugilidae fishes are subtropical species unknown to be breeding in or regularly migrating to regions north of Cape Cod (Ayvazian et al., 1992; Bigelow & Schroeder, 1953). Summer flounder is a temperate species more commonly found south of Cape Cod, but some individuals may make seasonal migrations to coastal Maine and are therefore not considered Gulf Stream Orphans (Bigelow & Schroeder, 1953).

### 3.2 Nearshore surface temperature anomaly

GAMs fit to average daily surface temperature data from the Portland Harbor tide gauge explained 95.9% of the deviance. Results indicated that 2014-2015 and 2017-2019 had average annual temperature anomalies below the CRP average (Fig. 3) These cooler years were characterized by sustained periods of daily temperatures beneath the modeled daily average in winter (December-February) and spring (March-May), and limited time above the modeled daily average in summer (June-August). In contrast, daily temperatures in both summers and winters of warmer years (2016, 2020-2024) typically remained above the modeled daily average. Notably, summer temperatures have been more than 1°C greater than modeled averages in each year since 2020 (Table 2).

**Table 2:**
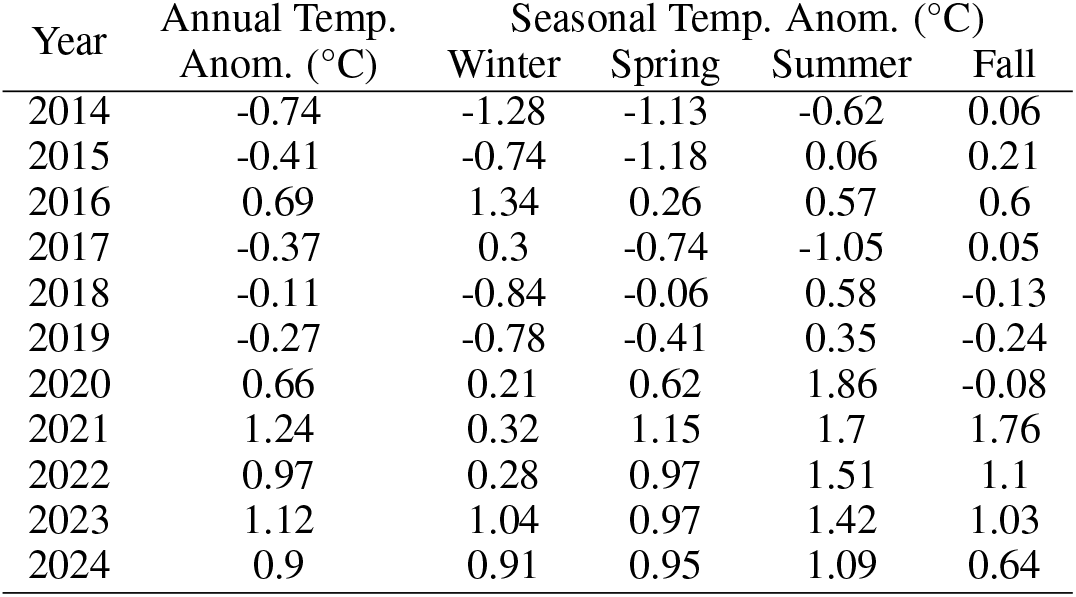
Portland Harbor tide gauge temperature anomalies presented as the difference between each years average temperature and the expected annual temperature as calculated from the 2003-2020 climate reference period (CRP). Anomalies are also presented at a seasonal scale, with winter referring to December-February, spring referring to March-May, summer referring to June-August, and fall referring to September-November. Negative values are cooler temperatures than expected compared to the CRP, and positive values are warmer temperatures than expected compared to the CRP.

**Figure 3:**
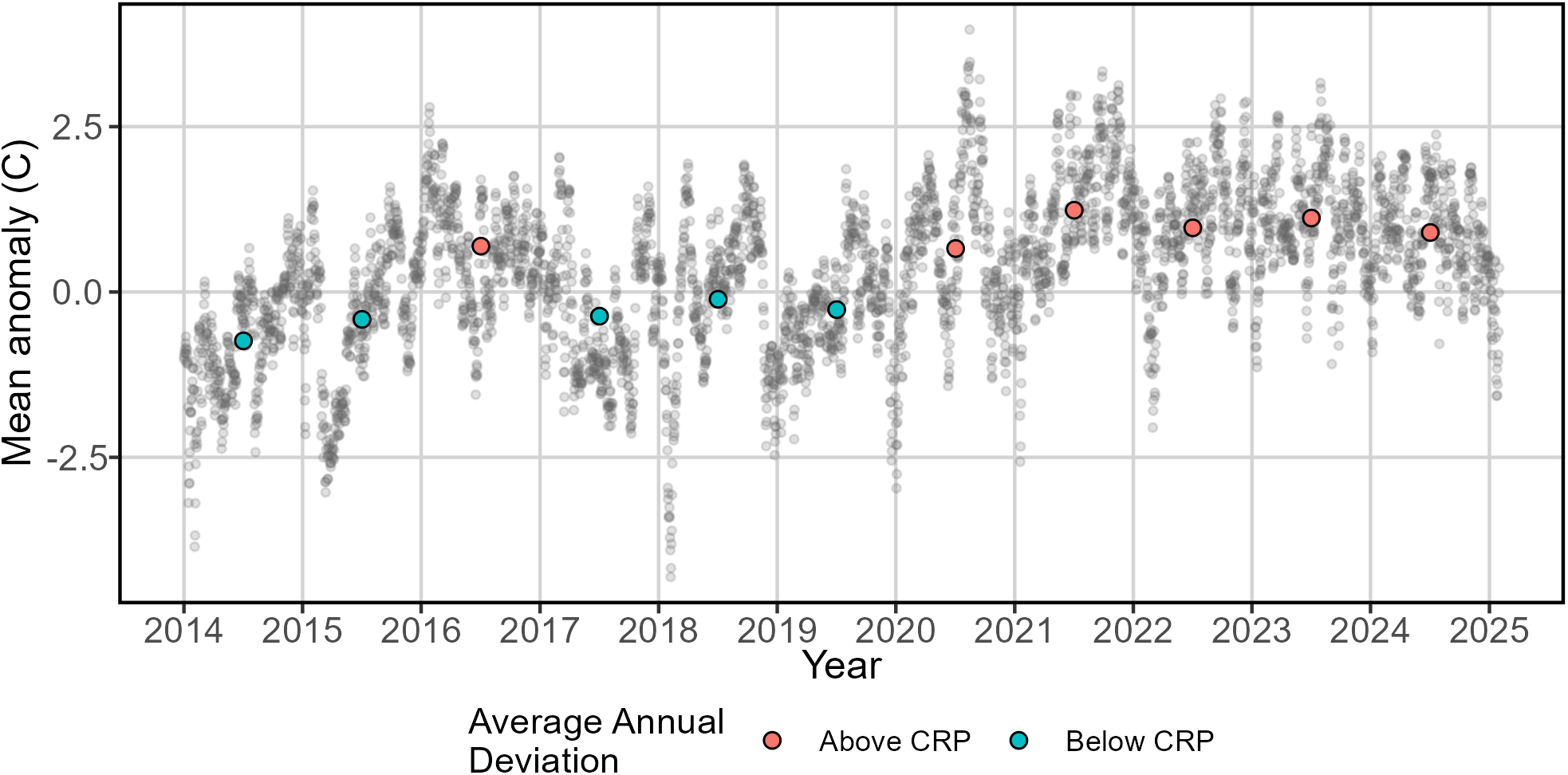
Temperature anomalies compared to the average daily temperatures estimated from the Portland Harbor Buoy 2003-2020 climate reference period. Gray dots indicate daily anomalies (in řC). Colored dots indicate average annual anomalies; red dots indicate anomalies above the CRP estimate, and blue dots indicate anomalies below the CRP estimate.

### 3.3 Weekly growth rates

The process of filtering length frequency distribution data to include only age-0 fish produced between 2 and 5 weeks of data per year to estimate herring weekly growth rates. Herring were caught in only the first week of sampling in 2023, so growth rates could not be estimated for that year. Herring growth rates in 2024 were estimated from data that had no sampling in weeks 32 to 38 because of vessel mechanical issues; we therefore had low confidence in the results for this year and did not include them in further analyses. The earliest week of age-0 herring detection was generally in week 25 (SD: 1.2 weeks), and the latest week of detection varied between week 26 and week 36 (mean: week 30, SD: 3.2 weeks). The wide range of the latest week of detection is partially caused by variability in sampling season end dates but is also caused by high interannual catch variability, decreasing herring catch through the sampling season, and infrequent encounters after the early summer period. Weekly growth rates for age-0 herring varied from 1.4 to 3.9 mm per week in warmer years and from 1.2 to 5.0 mm per week in cooler years (Fig. 4). There are many studies on larval herring growth rates in the Northwestern Atlantic, but few for juveniles with which we can compare our results. There is some evidence supporting growth rates of 2-2.5 mm per week for age-0 herring in the Bay of Fundy during the spring and early summer, but this is based on a single year of data collection (Das, 1972). Welchs t-tests indicated no significant difference when comparing mean growth rates between warmer and cooler years (t(4.907)=0.467, p=0.661).

**Figure 4:**
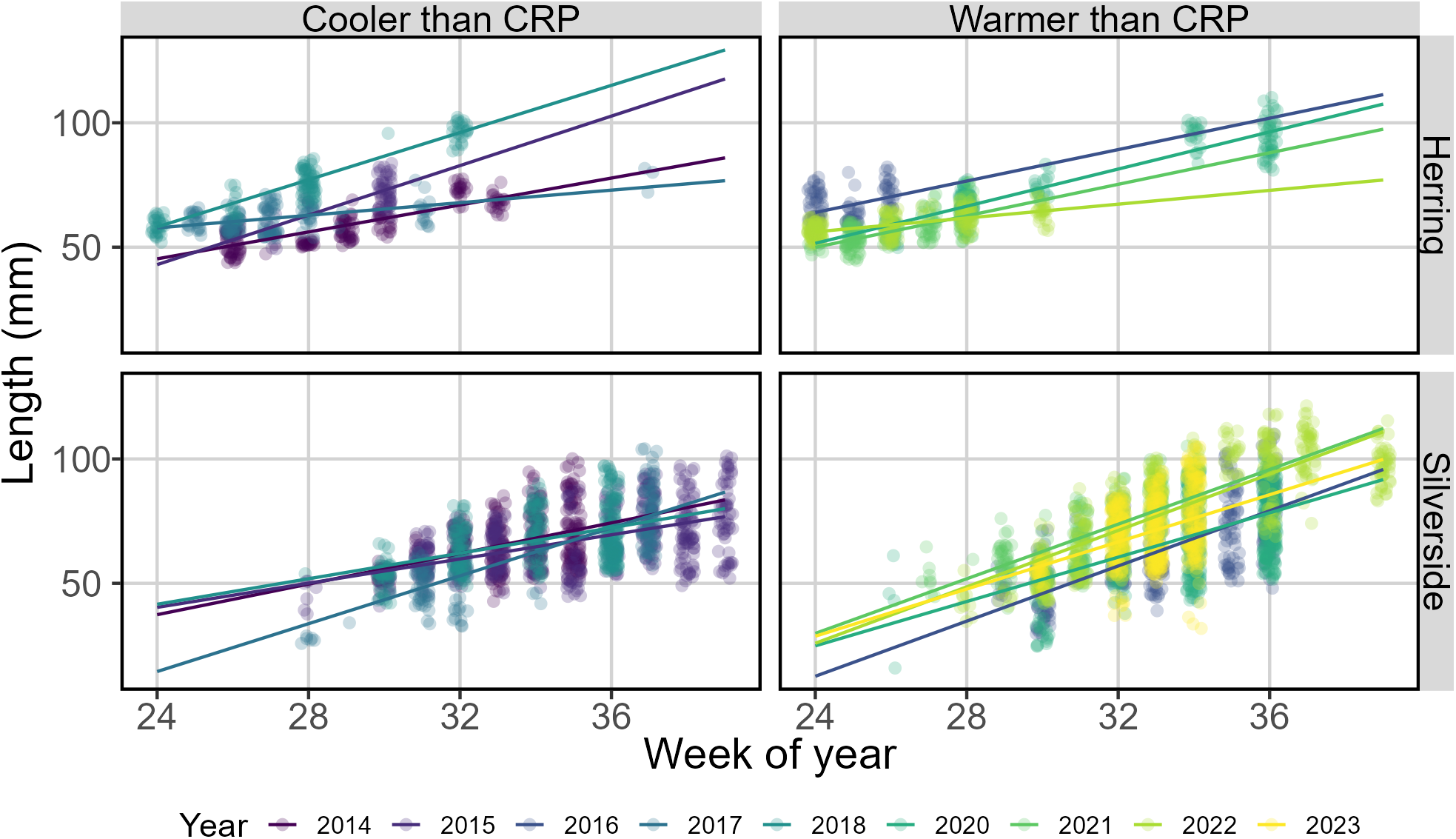
Weekly length distributions (in mm) of age-0 herring (top row) and silverside (bottom row) in years designated as having average annual temperature anomalies cooler (left column) and warmer (right column) than expected given the CRP. Lines indicate the calculated slope, or weekly growth rate. Colors of lines and dots indicate year.

Silversides were caught more consistently than herring, and between 2 and 8 weeks of length frequency data for age-0 individuals remained after data filtering. We excluded growth rates estimated for 2024 because of the gap in sampling data noted above. The earliest week of age-0 silverside detection was generally in week 29 (SD: 1.8 weeks), and the latest week of detection was usually the last week of the sampling season (mean: week 36, SD: 2.2 weeks). Silversides grew 4.5 to 6.2 mm per week in warmer years and 2.6 to 4.9 mm per week in cooler years (Fig. 4). Other studies have reported post-metamorphic age-0 silverside growth rates of around 5 mm per week for silverside populations in coastal New England (Barkman & Bengtson, 1987; Conover & Murawski, 1982; Pringle & Baumann, 2019), which is similar to our results. Welchs t-tests indicated growth rates were significantly different between warmer and cooler years (t(4.758)=-3.298, p=0.035), with a mean growth of 3.5 mm per week in cooler years and a mean growth of 5.3 mm per week in warmer years.

### 3.4 Effects on catch

Herring catch was significantly affected by the interaction of week and temperature, tidal height, surface temperature, and factor variables for bottom substrate and year (Fig. 5). Factor variable terms for weather and tidal stage and the random effect based on site were identified as not significant predictors and removed. The final model explained 56.7% of the deviance in catch. There was a strong effect of week on catch, as evidenced by the smooth representing the interaction of week and temperature. The strongest positive effect of week on catch occurred in weeks 30 and earlier and when temperatures were between 15 and 20°C. After week 30, there was an increasingly negative effect of week on catch regardless of temperature. The relationship between catch and tidal height was shaped like a sigmoidal curve, with a negative effect when the Portland Harbor tide gauge was lower than approximately 1.5m above MLLW. Above this point, increasing tidal height had a slowly increasing positive effect. The relationship between catch and temperature had a more complicated polynomial shape, but there was a clear positive effect when temperatures were between approximately 15 and 19°C. The effect of sand and sand/gravel substrate on catch was not significantly different, but there was a significant negative effect of mud substrate as compared to sand/gravel. All years except 2017 and 2020 had a significantly negative effect on catch as compared to 2014. The strongest negative effect was for 2023, which had the lowest mean herring catch per unit effort. Herring were caught in only the first week of sampling in this year. This low catch may be more reflective of a timing mismatch than a true decrease in abundance; sampling started later than usual and anecdotal observations indicated that herring had already used the nearshore region and left before the first seine operations were conducted.

**Figure 5:**
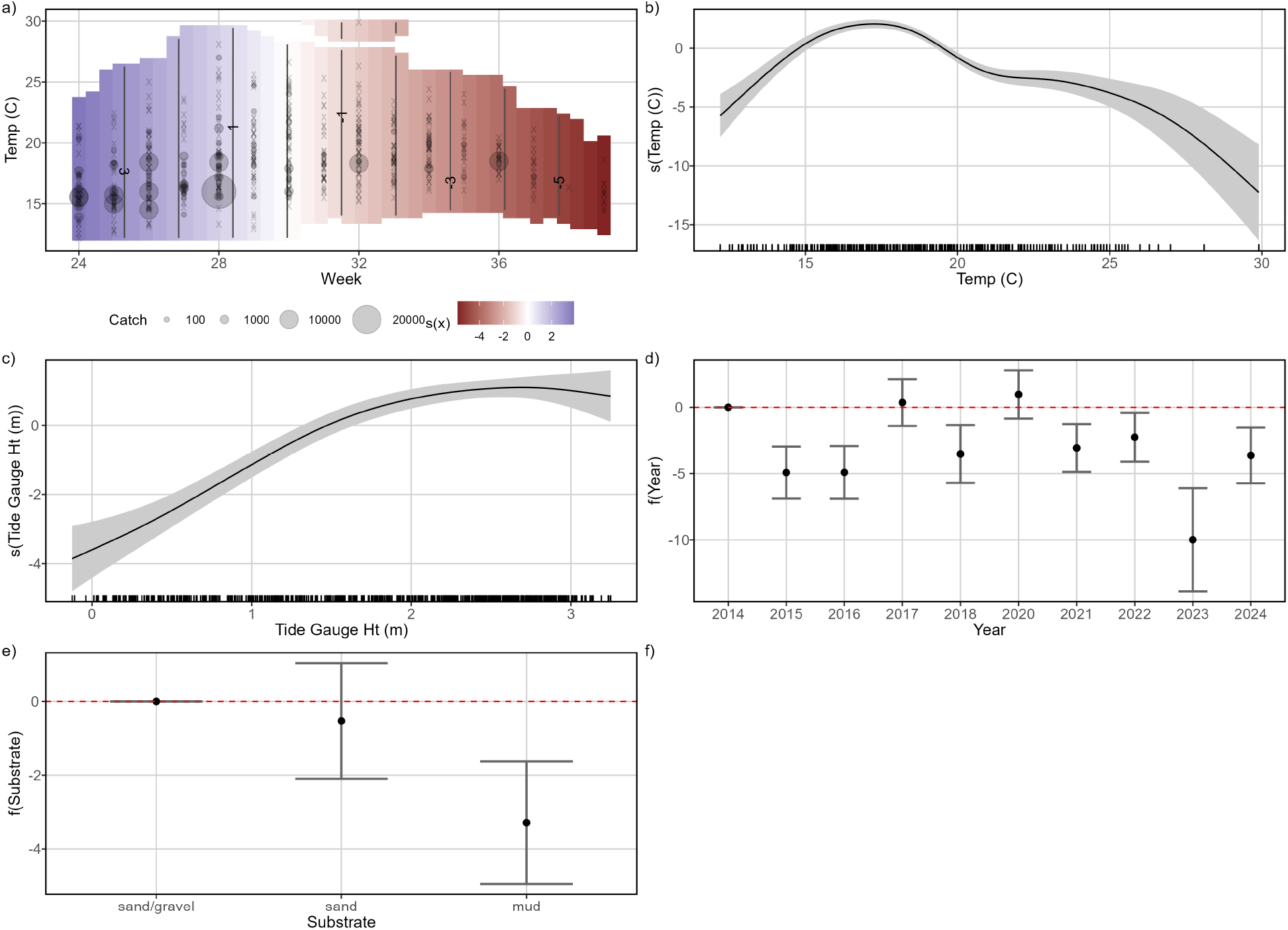
Smooth terms for the herring weekly catch GAM. Terms include (a) the interaction of week and temperature, (b) temperature, (c) tidal height (m) compared to MLLW, (d) factor effect of year, and (e) factor effect of substrate type.

Explanatory variables included in the final silverside catch model were the interaction of week and temperature, factor variables for weather conditions and year, and a random effect based on site (Fig. 6). Terms for temperature, tidal height, substrate type, and tidal stage were identified as not significant predictors of catch and removed. The final model explained 53.5% of the deviance in catch. In general, there was a positive relationship between catch and week, with the highest catches and a positive effect occurring in week 32 or later. However, the shift from week having a negative effect to week having a positive effect on catch could occur earlier in the summer (as early as week 30) if surface water temperatures were at or above approximately 18°C. The strongest negative effect on catch occurred in cool (>15°C) waters before week 30, and the strongest positive effect occurred in moderate temperatures (15-19°C) in weeks 33-38. Weather conditions also affected silverside catch. Compared to full sun, overcast conditions had a positive effect. There was no significant effect of partly cloudy or rainy conditions compared to full sun, though it should be noted that there were few samples taken in the rain. When holding all other variables fixed, the random effect of site had the highest variability in the sites near the mouth of the Presumpscot River (Audubon to Brothers North sites). For all years but 2015, there was a significant positive effect of year on catch as compared to 2014. The strongest positive effect was for 2020, which also had the highest mean silverside catch per unit effort.

**Figure 6:**
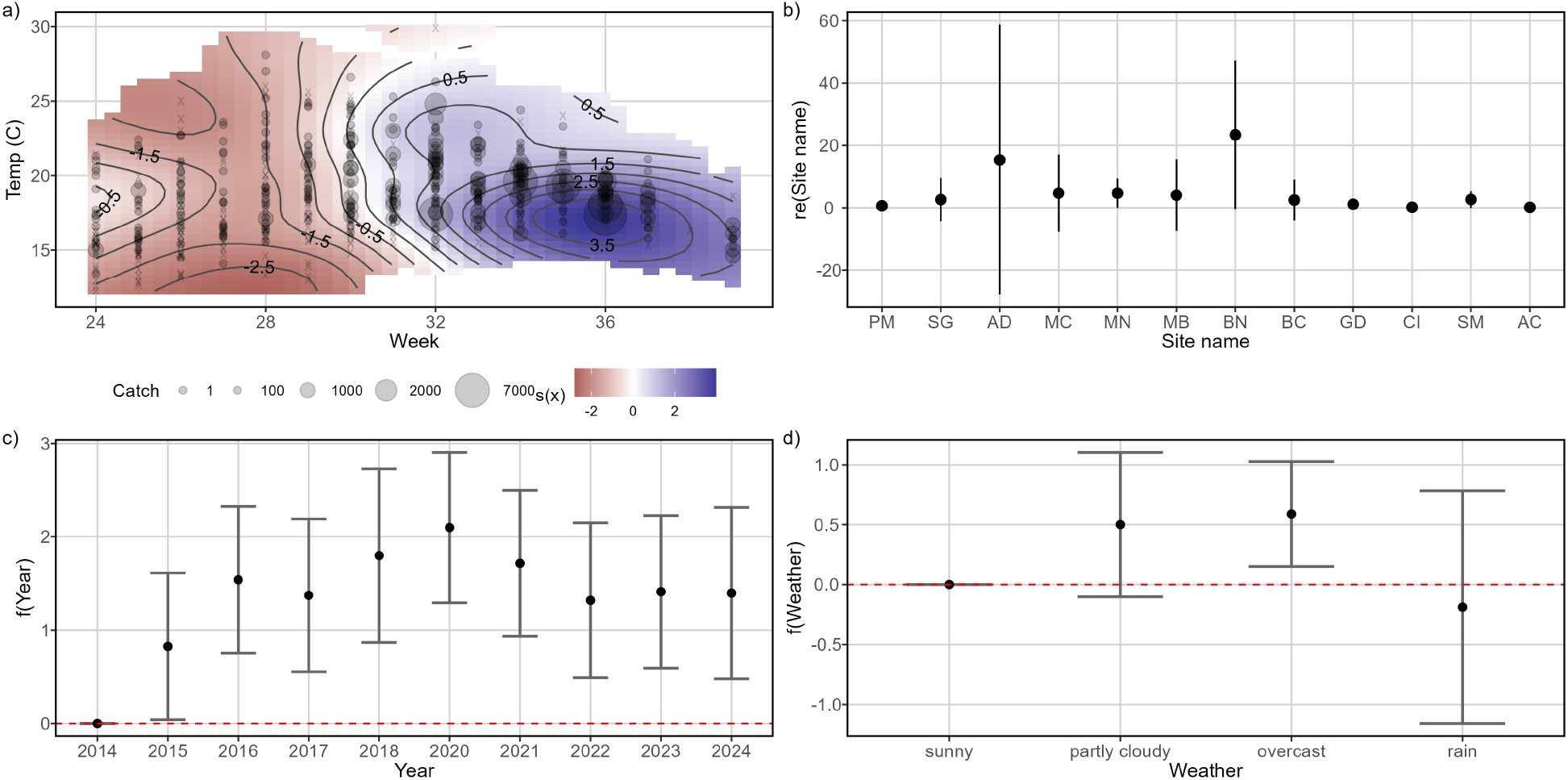
Smooth terms for the silverside weekly catch GAM. Terms include (a) the interaction of week and temperature, (b) random effect of site, (c) factor effect of year, and (d) factor effect of weather conditions.

### 3.5 Spatiotemporal community structure

NMDS analysis compressed site-specific seasonal community compositions to two ordination axes. The resulting stress coefficient was 0.13, indicating a sufficient goodness of fit and limited distortion. Fitting species as vectors within the ordination plot revealed that sand lance, herring, and silversides have the strongest associations with the ordination configuration (Fig. 7). Visualizing the total within sum of squares and average silhouette width for k in 1 to 20 clusters indicated the most support for k=3 clusters of similar community composition. ANOSIM confirmed a significant difference in community composition between these three clusters (r=0.761, p < 0.0001). The average CPUE for each species in each cluster was calculated and visualized, which revealed that cluster A typically had high herring catch, cluster B had generally low catch for all species, and cluster C had high silverside catch (Fig. 7).

**Figure 7:**
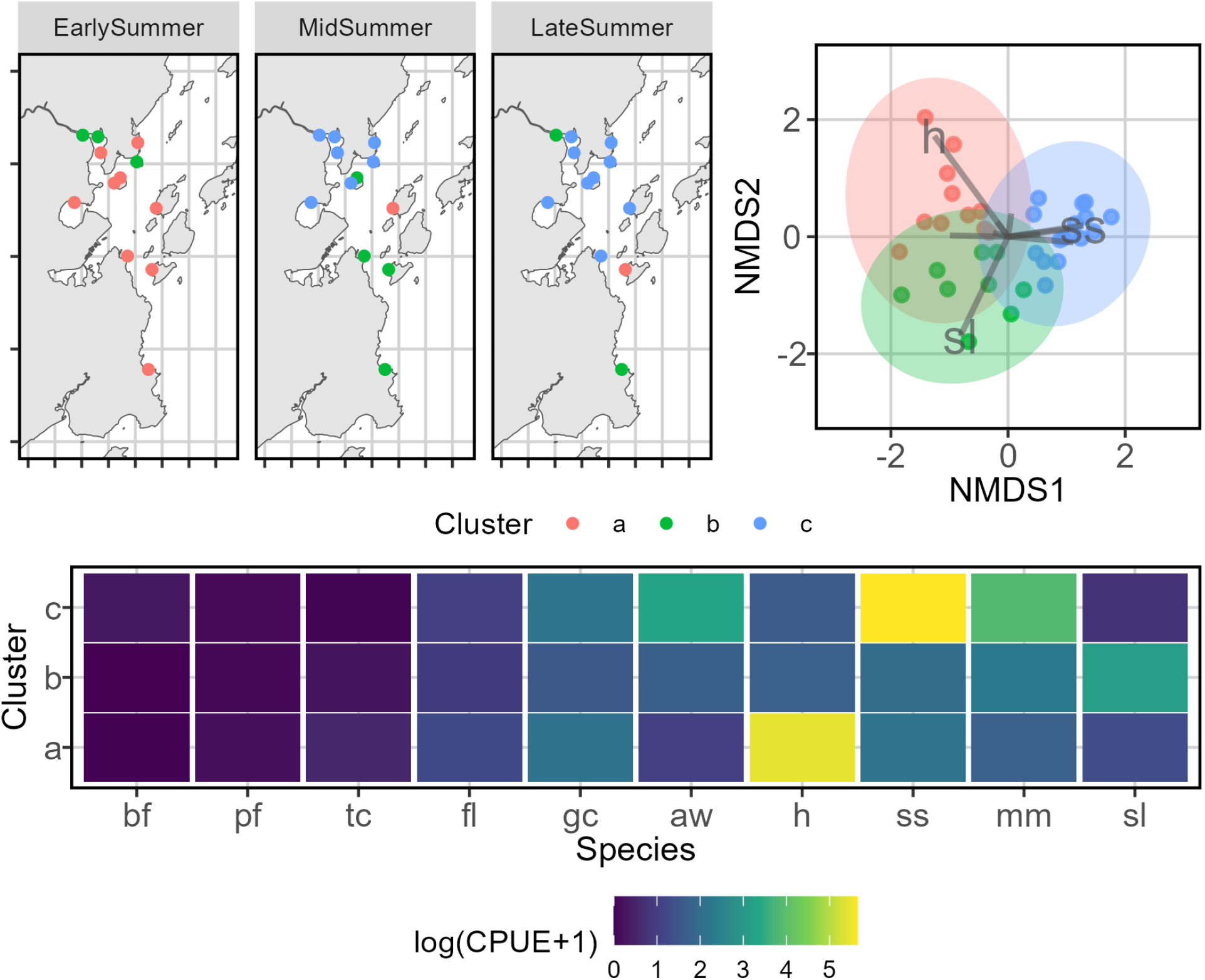
Nonmetric multidimensional scaling and k-means cluster analysis of community composition among all sampling sites. Clockwise from top left, the panels illustrate (a) the cluster of similar community composition found at each sampling site in the early summer (weeks 24-28 of each year), mid summer (weeks 29-33), and late summer (weeks 34-39) periods; (b) the NMDS ordination of each site-period combination (points) with coloring indicating the same clustering structure and herring (h), silverside (ss), and sand lance (sl) association to the ordination structure illustrated as vectors; and (c) the log-scaled average catch per unit effort (CPUE) for all species by cluster. Species are bluefish (bf), northern pipefish (pf), Atlantic tomcod (tc), winter flounder (fl), green crab (gc), river herring (aw), Atlantic herring (h), Atlantic silverside (ss), mummichog (mm), and American sand lance (sl).

SIMPER analysis across clusters provided quantitative confirmation of this visual analysis. When comparing the composition of cluster A to cluster B, herring (p=0.001, Contribution=62.5%), sand lance (p=0.007, Contribution=12.4%), and green crabs (p=0.016, Contribution=3.9%) combined to contribute more than 90% of the dissimilarity. Winter flounder, Atlantic tomcod, and northern pipefish contributed smaller, but still significant, amounts of dissimilarity. Cluster A had higher average CPUE for all species identified as significantly contributing to dissimilarity except sand lance, which had higher average CPUE in cluster B. When comparing clusters A and C, only catch of silverside (p=0.002, Contribution=49.6%) and herring (p=0.017, Contribution=31.0%) had significant contributions to dissimilarity. Herring had a higher average CPUE in cluster A, and silverside had a higher average CPUE in cluster C. Dissimilarity between clusters B and C was driven by catch of silverside (p=0.001, Contribution=66.2%), mummichog (p=0.025, Contribution=12.4%), and bluefish (p=0.031, Contribution=0.09%). All three of these species had higher average CPUE in cluster C as compared to cluster B.

When plotted across space and time, divergent spatiotemporal patterns of habitat use can be identified (Fig. 7). In the early summer period, most sampling sites (9 of 12) are best described by cluster A. Only one site belongs to cluster A in the mid- or late summer periods, which matches the noted trend in decreasing herring catch through the summer. Cluster B is present across all three seasonal periods. The mixed-composition community structure of this cluster best describes sites with low CPUE for all species included in the modeling efforts. Sand lance has the highest CPUE of all species within this cluster, but it should be noted that sand lance catch was uncommon and mostly observed in the late summer period at the two southernmost sites. In early summer period, but many sites were best described by this cluster in the mid-summer (7 sites) and late summer (9 sites) periods. This matches the noted trend in increasing silverside catch through the summer.

NMDS ordination of community composition aggregated across all sampling sites per year also had sufficient goodness of fit across two ordination axes (stress=0.08). Herring, silverside, and northern pipefish had the strongest associations with ordination configuration, and k-means clustering analysis supported the selection of two clusters (Fig. 8). ANOSIM confirmed a significant difference in community composition between clusters (r=0.81, p=0.021). Visual representations of ordination plots and average CPUE per species within both clusters indicated that cluster A was characterized by high silverside catch and cluster B was characterized by high herring catch (Fig. 8). SIMPER results highlighted herring as the only species contributing a statistically significant amount of the dissimilarity between the clusters (p=0.001, Contribution=67.6%). Casco Bay community composition in 2014 and 2017 was best characterized by cluster B, and all remaining years were best characterized by cluster A. Though there was no clear connection between average annual temperature anomaly and community composition cluster, it should be noted that both 2014 and 2017 were cooler than the CRP and had extended periods in the spring and summer with below-average daily temperatures.

**Figure 8:**
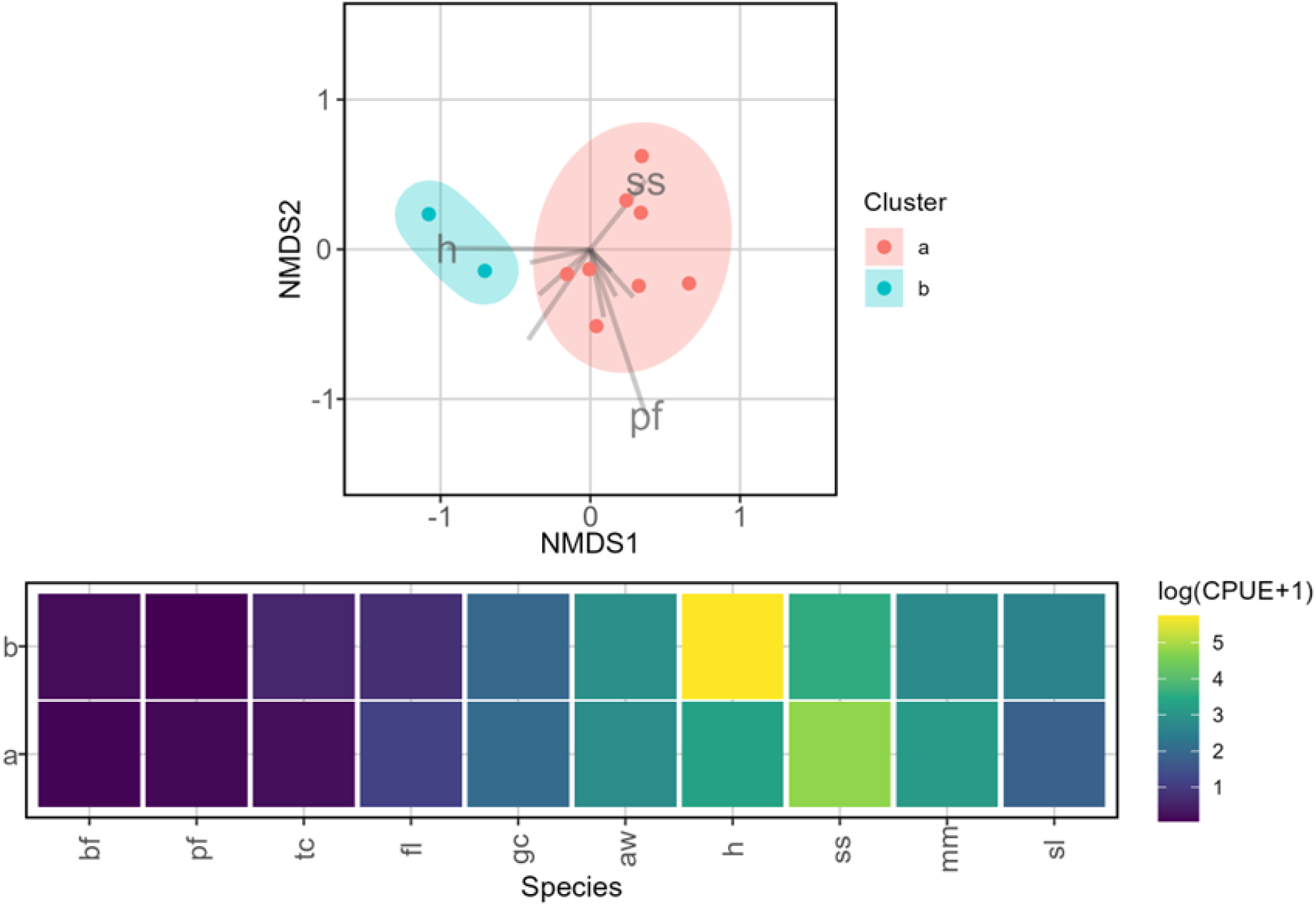
Nonmetric multidimensional scaling and k-means cluster analysis of community composition among all sampling sites. The top panel illustrates the NMDS ordination of each annual Casco Bay-wide annual community compositions, with the color of the points indicating the cluster and herring (h), silverside (ss), and northern pipefish (pf) association to the ordination structure illustrated as vectors. The top panel illustrates the log-scaled average catch per unit effort (CPUE) for all species by cluster. Species are bluefish (bf), northern pipefish (pf), Atlantic tomcod (tc), winter flounder (fl), green crab (gc), river herring (aw), Atlantic herring (h), Atlantic silverside (ss), mummichog (mm), and American sand lance (sl).

## 4 Discussion

Many studies have identified responses to hydroclimatic changes at a species-specific level (Le Bris et al., 2018; Pershing et al., 2015), and others have focused on responses at higher levels of ecological organization and at large spatial scales like the Northeast US Continental Shelf Large Marine Ecosystem (Fenwick et al., 2024; Lucey & Nye, 2010). Fewer studies have explored temperature-linked changes to the community ecology of GoM nearshore regions. We were interested in characterizing seasonal and interannual patterns of Casco Bay species assemblages, with an emphasis on the effect temperature has on these patterns. Many ecologically and economically important species use this area, connecting it to larger spatial scales and several levels of biological organization. Atlantic silverside and Atlantic herring, as highly abundant and important forage species with differing adaptability to temperatures experienced in Casco Bay, were used as focal species to estimate the effects of temperature on growth and abundance.

The phenology and relative abundances of Casco Bay nearshore fishes, and the effects of increasing nearshore surface temperatures, were illustrated through the analysis of 11 years of seine survey data. Catch was dominated by a few species of pelagic fish, including herring and silverside. There was a clear seasonal progression to community structure, which is closely tied to the relative abundances of these focal species. Most sites in the early summer period (weeks 24-28) were best described by high herring catches and relatively low catches of other species.

Similarly, GAMs indicated a strong positive effect of both early weeks and cooler temperatures on herring abundance, but a strong negative effect for the same conditions on silverside abundance. The mid-summer period (weeks 29-33) is a transitional period, where temperatures rapidly increase and community assemblage shifts to generally be more silverside-dominated as YOY silverside recruit to the seine and herring leave the nearshore area. Note that we have no information on how far or deep juvenile herring go when then leave the reach of our beach seine, though concurrent collection of environmental DNA (eDNA) samples may shed light on fine-scale herring spatiotemporal distributions. This is especially true of sites in and around the Presumpscot River estuary, which are typically warmer than the sites with less freshwater influence. The late summer period (weeks 34-39) had a strong negative effect on herring regardless of temperature. Though there was generally a strong positive effect on silverside abundance in this period, this effect was weakened when surface temperatures exceeded 20°C.

The ties between community structure and seasonal warming were evident when analyzing at site-specific and short-time scales. We hypothesized that longer-term warming would also result in changes in community structure and somatic growth at larger spatial and temporal scales. It was expected that years with average temperatures exceeding the CRP average would benefit silverside and disadvantage herring, resulting in altered community structure and opposing effects on growth. Community structure viewed through an annual, whole-Bay lens did not clearly follow this pattern. Though two years categorized as cooler than the CRP average (2014 and 2017) had relatively high herring abundance and were best described by a herring-dominated community structure, the remaining cool years (2015 and 2018) had relatively low herring abundance and were best described by a silverside-dominated community structure. All years categorized as warmer were best described by a silverside-dominated community structure.

Interestingly, some of the lowest estimated herring weekly growth rates occurred in years with high herring relative abundance (2014 and 2017), while years with low herring relative abundance (2015 and 2018) had the two highest estimated growth rates. This may indicate a density-dependent effect on growth, which has previously been noted in GoM herring populations (Becker et al., 2020), although this was for adult herring. Though there was a trend of decreased herring weekly growth rates in warmer years, this trend was not significant. Population dynamics of herring have notably high interannual variability, with a wide range of biotic and abiotic factors and selective processes acting on recruitment, growth, distribution patterns, and mortality (BeaudrySylvestre et al., 2024; Becker et al., 2020; Burbank et al., 2023). The characteristics of the population observed in nearshore Casco Bay seines are likely influenced by processes at much larger spatial and temporal scales than captured in this study.

Silverside weekly growth rates were significantly higher in warmer years than cooler years. Silverside relative abundance increased over the 11-year study period, which may also be related to the increasing summer nearshore temperatures experienced in the study area over this time. The short lifespan, early summer spawning, and nearshore distribution of the YOY silverside primarily caught in our seine surveys closely links their population dynamics to short-term, local processes. This, combined with their highly abundant and ubiquitous nature, may make them a useful indicator species for coarse-scale structure and functioning of the Casco Bay ecosystem.

Reduced relative abundance of certain forage species in nearshore GoM ecosystems, including herring, could negatively impact nearshore trophic dynamics. Changes in prey size, distribution, relative abundance, and community structure will impact piscivorous predators that have evolved to exploit nearshore forage fishes (Ball et al., 2007; Falke et al., 2024; A. T. McGowan et al., 2022; D. W. McGowan et al., 2019). Our study has illustrated temperature-related changes in seasonal herring nearshore habitat use and indicates that long-term temperature increases could alter temporal patterns of distribution in nearshore GoM areas. Though silverside may grow faster and increase in abundance with warmer temperatures, they may not fill the same role in trophic dynamics as herring. In the early summer, silverside in the nearshore region were much larger than herring. This size difference could preclude silverside from being a prey item for juvenile piscivorous fishes using the nearshore region as a nursery area. Prey size is an important limiting factor in the ontogeny of piscivory in striped bass and bluefish, both of which are important sportfish and predators in the GoM (Scharf et al., 2009). Herring and silverside also have differing spatial distribution patterns; silverside typically inhabit extremely shallow nearshore regions (Conover & Murawski, 1982), while herring are more widely distributed (Boyar, 1968; Creaser & Libby, 1986, 1988). If biomass of forage fishes shifts closer to shore and into shallow areas, it may alter prey availability to larger-bodied predators or force changes in predator distributions.

The dynamics of shallow nearshore ecosystems and the forage species that use them are relatively understudied (Lankowicz et al., 2020; A. T. McGowan et al., 2022). The abundances and distributions of taxonomic groups in these highly heterogeneous areas are spatially and temporally variable along environmental gradients. Some environmental gradients may directly influence the behavior of a species (temperature, dissolved oxygen, salinity); this may modify interspecific interactions (competition, predation) and lead to further indirect effects. Identifying the environmental patterns that drive fish distribution and community structure could provide predictive insight under alternate climate scenarios, as well as for much larger spatial scales and biological organization levels.

The results of this study indicate that increasing surface water temperatures will likely lead to altered timing of nearshore habitat use by Atlantic herring, which may affect their population dynamics. Further exploration into interannual variation in juvenile herring population dynamics and individual growth may add context to the steep decline in recruitment and associated reduction in spawning stock biomass seen since 2012, which has persisted despite reduced fishing pressure (NOAA, 2024). This produces natural follow-up questions regarding altered interspecific interactions within nearshore ecosystems, trends in average energetic content of herring as compared to silverside, as well as the potential of interspecies competition between herring and silverside for prey in the nearshore regions. Though answering these questions is beyond the scope of this paper, it is clear that current Casco Bay nearshore temperatures are still within the thermal tolerances of Atlantic silverside, which has led to increased abundance and faster growth. Monitoring efforts like the Casco Bay Aquatic Systems Survey should be continued, and possibly expanded in temporal scope, to further explore the effects of shifting community structure on the trophic ecology of nearshore regions.

## Funding and Acknowledgements

The Casco Bay Aquatic Systems Survey (CBASS) was launched in 2013 with a gift from an anonymous donor and has received generous support in recent years from the Scanlan Family Foundation. The authors would like to thank Isabelle Sée, Tait Nygaard, and Alec Bollinger at the Quahog Bay Conservancy (QBC) for assistance in data collection, as well as the numerous REU students and staff members at GMRI and summer interns at QBC who have supported CBASS over the past 11 years.

## Author contributions

Conceptualization: GS, KL

Data curation: ZW, AW, SB, KL

Formal analysis: KL

Funding acquisition: GS

Methodology: KL

Supervision: GS

Visualization: KL

Writing - original draft: KL, CS

Writing - review & editing: KL, GS, CS, ZW, AW

## Data availability

Data will be made availablre upon request to the corresponding author.

